# The Pulfrich Effect in Virtual Reality

**DOI:** 10.1101/2024.03.21.585956

**Authors:** Anthony Loprete, Arthur G. Shapiro

**Author notes:** Authors’ addresses: Anthony LoPrete, Bioengineering Graduate Group, University of Pennsylvania, Philadelphia, Pennsylvania, USA, 19104-4202; Arthur G. Shapiro, Center for Neuroscience and Behavior, American University, Washington, D.C., USA, 20016.

## Abstract

The Pulfrich effect, a visual phenomenon where a neural delay in one eye produces a depth misperception, has been directly studied on flat-panel displays but not in virtual reality (VR) environments. Through a series of three experiments, we investigated the relationship between luminance, contrast, dot spacing, and optical blur on the Pulfrich effect in VR and on the perception of motion. In the first two experiments, we found that low-reflectance stimuli produce a stronger Pulfrich effect than high-reflectance stimuli in VR, a result further accentuated by background luminance. Furthermore, the primary experiment showed that nullifying helix rotation motion is a powerful way to study the magnitude of the Pulfrich effect. With data from the first experiment, we developed a compelling VR illusion in which changing the color of a helix reverses the direction of perceived motion. Experiment 2 elaborated that low-reflectance stimuli only produce a stronger Pulfrich effect than high-reflectance stimuli when the stimulus is moving away from the delayed eye. Data from the first two experiments were successfully captured by power law function fits and linearized by plotting Pulfrich effect strength against logit-Michelson contrast. Our third experiment revealed that increasing blur and dot count in the helix stimulus increased the likelihood of perceiving up or down motion. All three experiments in tandem show that investigating well-known visual illusions such as the Pulfrich effect in virtual reality has the potential to reveal insights into visual perception as well as inform us about the effects of contrast and asymmetric lighting in spatial computing.

CCS Concepts: • **Computing methodologies** → **Perception**; **Virtual reality**.

## 1 INTRODUCTION

VR and AR displays are becoming increasingly common across science, industry, and entertainment. In vision science, many experiments now present stimuli in VR on head-mounted displays (HMDs) as a means to study perception under more naturalistic viewing conditions [13, 32]. Among other topics, VR has already been used to study interpersonal space [51], 3D motion perception and feedback [8], distance perception [17], scene saliency [44], proprioception with hand-tracking [16], and color awareness during naturalistic scene viewing [6]. VR technologies work by creating images that simulate the light array typically provided by the natural environment [35]. The brain extracts depth information (often referred to as “depth cues”) from the light array, as it would typically for natural scenes. The multiple sources of depth information in natural scenes often provide a consistent, statistically reliable story about the external world.

One benefit to VR is that, like other laboratory displays such as haploscopes, the technology can present depth cues that provide conflicting information; for instance, an object’s spatial disparity (differences in perceived position for each eye) could indicate it is moving away from the observer, while its increasing relative size could indicate it is moving closer. Such conflicts between stimuli have been used to create new visual phenomena, give insights into underlying visual processes, and allow for investigations of cue combination [37, 42]. For instance, the contrast asynchrony illusion [40, 41] juxtaposes luminance information modulated synchronously and contrast information modulated asynchronously; the double drift illusion juxtaposes global motion with internal spin [12, 21, 23, 39, 49]. Much can be learned from how and in what conditions the visual system resolves conflict: for instance, in the contrast asynchrony illusion, the perceptual system maintains a separation of luminance and contrast, whereas, in the double drift illusion, global and internal motion are kept separate in the fovea but combine in the periphery. Both these resolutions are a different solution to what Anne Treisman referred to as the binding problem: given that the visual system encodes different sources of spatial (and, in our case, depth) information, how does the visual system weigh this information to create an understanding of the visual world? [3, 48]

Here we use a VR system to place two different types of depth cues in conflict with each other: the Pulfrich effect and the Hess effect. The Pulfrich effect [30] is a phenomenon where an interocular delay (often produced by a dark filter over one eye) introduces depth from disparity (a binocular depth cue). A typical demonstration of the Pulfrich effect is the Pulfrich pendulum. When the swinging pendulum of a grandfather clock is observed with a dark filter placed in front of one eye under binocular viewing conditions, the pendulum is perceived as following an elliptical trajectory. The accepted explanation for this effect is that the filter produces a neural delay between the eyes. This interocular delay causes the motion to follow a new 3D trajectory consistent with the binocular geometry. Interest in the Pulfrich effect as a tool in vision science stems from its ability to study interocular delays induced by low-level stimulus features as correlates with perceived depth or motion [4]. Previous work has also shown similar effects from interocular differences in contrast [31] and spatial frequency [3].

The Hess effect [14, 52] is a phenomenon where changes in luminance contrast (a monocular depth cue) between the background and stimulus influence depth perception due to processing speed discrepancies between low and high contrast stimuli. In the Hess effect, the visual system responds faster to moving targets with higher contrast than to moving targets with lower contrast. Stereopsis, which is affected by luminance contrast, is also influenced by the Hess effect [18, 36].

We juxtaposed the Pulfrich and Hess effects by using a double-helix hybrid-motion stimulus [43]. In this stimulus, two columns of dots oscillate horizontally in position with opposite phases. The double helix hybrid-motion stimulus encodes two different types of motion signals at different spatial scales: left-right motion at high spatial frequencies, and up/down motion at low spatial frequencies [37]. Hybrid motion illusions can be interpreted as video extensions of hybrid image illusions [29]. Two examples of the double-helix stimulus are shown in Figure 1. Panel (A) of Figure 1 is the double-helix stimulus implementation of the Hess effect. The two vertical arrays of dots still oscillate horizontally in both luminance and position on the same period; depending on the phase of the dot’s luminance oscillation, perception of helical rotation will be perceived as front-left or front-right. Panel (B) of Figure 1 depicts the hybrid motion component of the double helix stimulus. In the left helix, motion is generally perceived as left/right. When the stimulus is blurred (removing low spatial frequency information), motion is instead perceived as up/down.

**Fig. 1.**
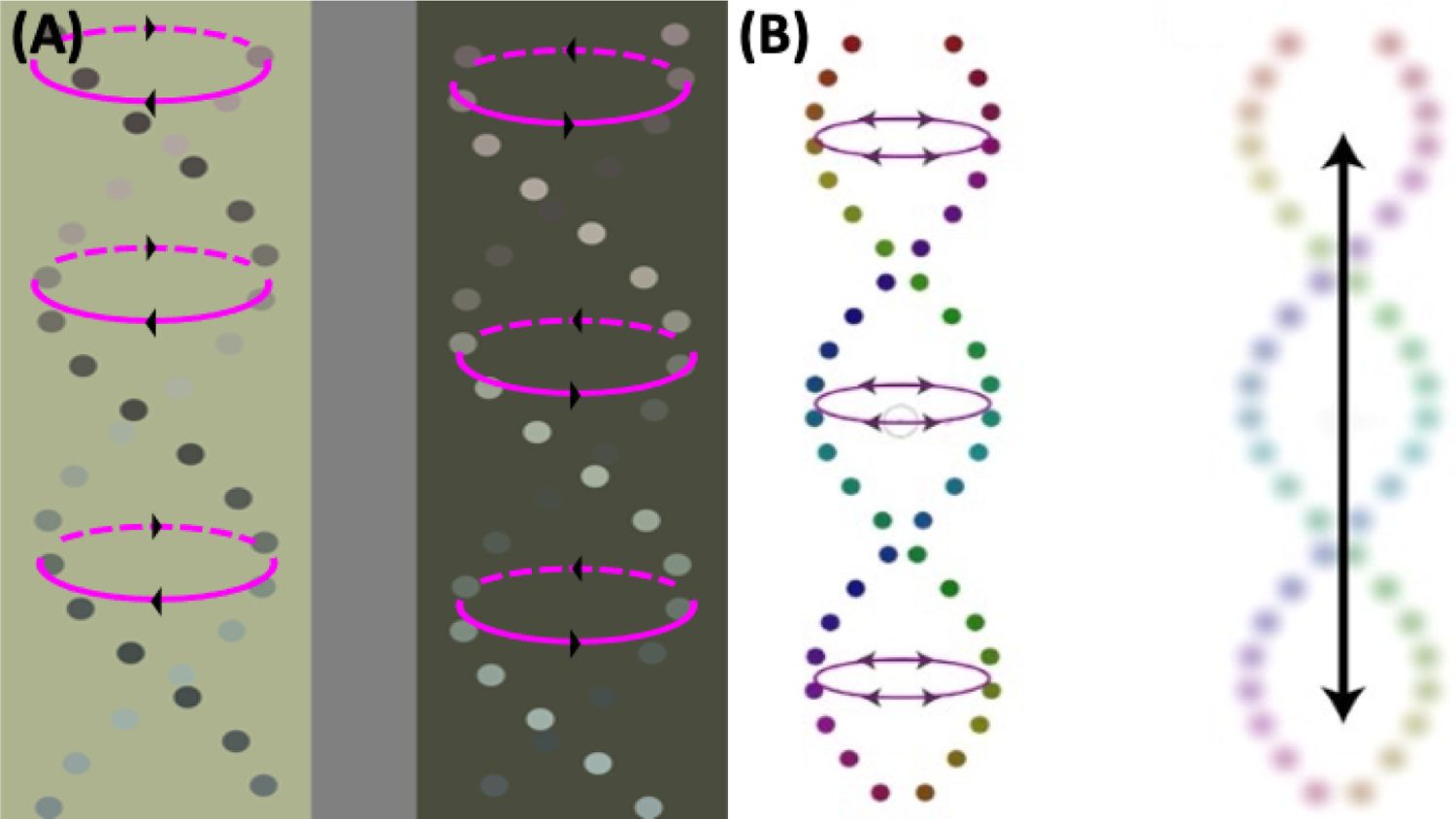
Two depictions of the helix rotation stimulus that we used for the main experiment in this paper. In both panels, each dot moves horizontally across the display. **(A)** Helix rotation version of the Hess effect. This demo has been previously shown, first at the 2019 International Colour Vision Society meeting. High-contrast dots are processed more quickly than low-contrast dots. As a result, the left helix appears to rotate with the “closer” dots moving towards the left (“front-left”). Similarly, the right helix appears to rotate front-right. **(B)** Helix rotation version of hybrid motion. This panel first appeared in [43]. In the unblurred display, helix motion is perceived as side-to-side. When the display is blurred, perceived motion switches to up/down. In this sense, the helix encodes two types of motion. Therefore, perceived motion in the helix can be used as a measure of another variable. In this case, perceived helix motion can be used to measure image blur.

The purpose of the Pulfrich helix stimulus was to measure the contributions of local contrast and global luminance on the strength of the Pulfrich effect in VR. We re-reported the finding that lower luminance conditions correspond to greater Pulfrich effect strength [24]. However, we also revealed that this effect is greater for low-reflectance stimuli than for high-reflectance stimuli with the conditions of our experiment. We used this conflict between Pulfrich and Hess to create a new VR visual phenomenon in which stimulus reflectance unexpectedly led to massive changes in perceived motion. Additionally, our second experiment demonstrated that the stimulus-reflectance dependency on the Pulfrich effect in VR was driven by motion away from the filtered (delayed) eye. Lastly, we investigated the effects of low-frequency masking on hybrid-motion stimuli by performing a third study involving the helix rotation stimulus on a conventional flat-panel display with optical blur. The results of these experiments may prove useful when thinking about how to implement depth cues in VR environments.

## 2 EXPERIMENT 1: HELIX ROTATION IN VIRTUAL REALITY

In the first experiment, we addressed how the perception of depth in virtual reality depends upon the luminance of an object, interocular luminance differences (Pulfrich effect), and the relative luminance between objects and the background (Hess effect). To this end, we measured how luminance (white stimuli and black stimuli) and contrast (different brightness levels of a background) contribute to the perceived strength of the Pulfrich effect in VR with a nulling procedure. We also hypothesized that the direction of perceived motion in the helix stimulus would switch depending on the relative magnitude of the stimuli driving the Pulfrich and Hess effects.

### 2.1 Observers

Three observers participated in this experiment: the two authors and a third female naive observer. Observers had normal or corrected vision and no history of colorblindness. Experiments were conducted on the American University campus in accordance with the American University Institutional Review Board.

### 2.2 Equipment

The experiments were conducted with an HTC Vive Virtual Reality system. Observers were seated in a chair and instructed not to move their heads. Luminance parameters of the VR display were measured with a Photoresearch Spectroscan 670 spectroradiometer. The spectroradiometer was pointed at a screen of the VR display (in a manner identical to that shown in Figure 1 of [9]), from which we determined the gamma curve by measuring the luminance corresponding to several machine unit values spanning from 0 to 1. The maximum luminance of the display is 180.20 candelas per square meter (cd/m^2^) and the minimum is 0.28 cd/m^2^. Observers made inputs with the HTC Vive controllers.

### 2.3 Stimulus

The stimulus (depicted in Figure 2) was designed and rendered in the Unity-3D game engine. The stimulus was a 50-dot double-helix (two columns of 25 dots each). Each “dot” of the helix was a 1 cm-radius, solid-color, moderately lustrous sphere from Unity’s primitive objects. The stimulus was rendered in front of observers at an average distance of 0.69 meters, which was variable with head size and posture. The dots were equally spaced along the y-axis, and the helix location was constant in the virtual environment. The helix had a fixation dot located in the center. The dots oscillated laterally across the x-axis with an amplitude of 2.5 cm and an angular frequency of 4 rad/sec (2/π Hz). The temporal phase between the dots determined the overall direction of y-axis motion (up or down.) The virtual environment was illuminated with both Unity ambient lighting and an overhead lighting source. Small specular effects were exhibited by the helix dots due to the rendering parameters, as seen in Figure 2’s stimulus panel.

**Fig. 2.**
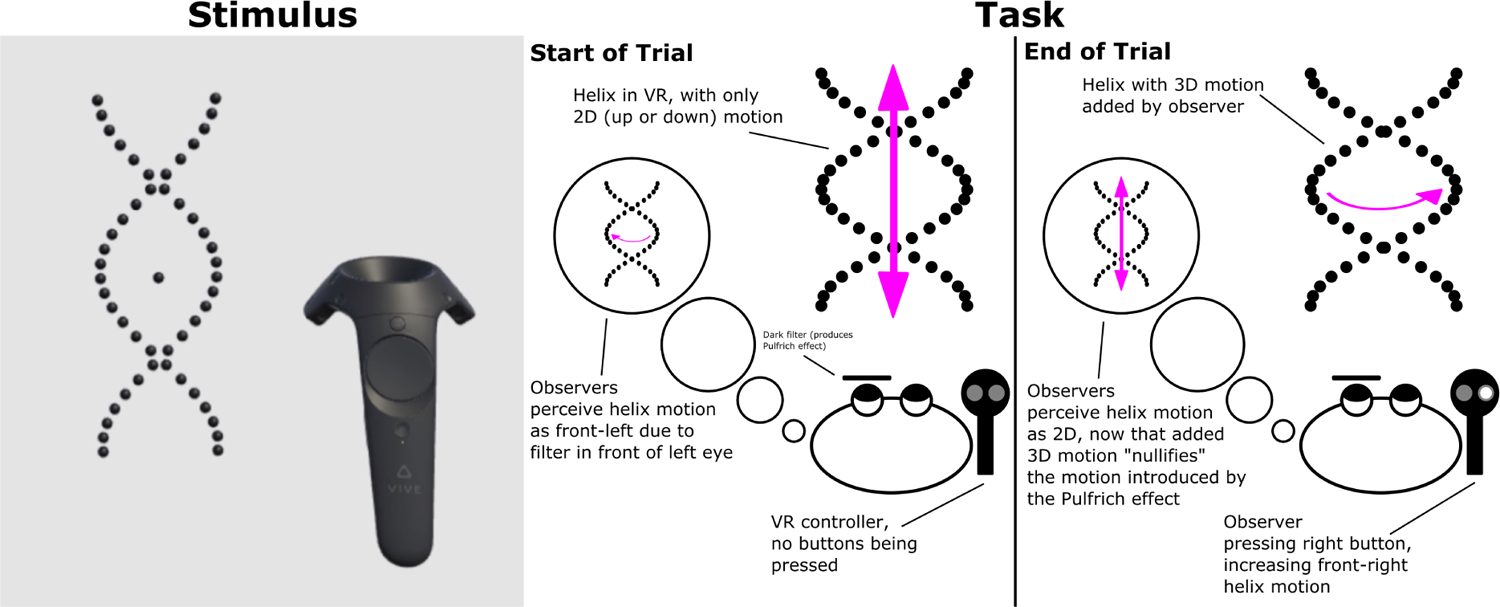
Stimulus: One trial from a session of the helix rotation experiment in Virtual Reality. Two stacked columns of dots rotate horizontally in a fronto-parallel plane. One eye’s screen on the VR display was darker than the other, causing the stimulus to appear as a three-dimensional rotating helix due to the Pulfrich effect. The controller for the Vive VR headset was rendered during the experiment. The image depicts a trial with dark dots in front of a bright background. The reflectance of the dots and the background varied parametrically. **Task:** Illustration of the task structure for experiment one. The Pulfrich effect makes dots moving in the fronto-horizontal plane appear to orbit in three dimensions. The observer nulls the depth effect by introducing rendered 3D motion in the opposite direction. When the motion due to the Pulfrich effect is fully canceled out by the 3D motion in the helix, the observer will perceive the motion as up or down 2D motion. The observer then confirms their response with a button press, and the next trial begins.

The stimulus backgrounds were 0.125, 0.3125, 0.5, 0.6875, and 0.875 in machine units (in RGB, the first background is (0.125, 0.125, 0.125), the second is (0.3125, 0.3125, 0.3125), etc.). These values correspond to 1.57, 12.95, 38.61, 78.55, and 132.77 cd/m^2^. These values were selected because they evenly sample the range of machine unit values between the maximum and minimum display values (which are used for the stimuli) given the number of background levels we were interested in testing (5) per session. In each trial, one eye of the VR display would be rendered behind a “filter.” This filter was a solid-black Unity canvas game object with an alpha value of 230 on an RGB 0-255 scale which covered the entire field of view for that eye only. The stimulus background was rendered as a solid color from Unity’s camera objects. The controller was rendered in the VR environment during experiments.

### 2.4 Experimental Procedure

The task for this experiment is depicted on the right of Figure 2. In a given session, the stimulus was rendered against 5 different backgrounds of different luminance values, and the dots that make up the helix would be rendered minimum or full Unity-specified reflectance (corresponding roughly to “black” and “white” dots, respectively), combining for a total of 100 trials per session, with each combination of stimulus parameters presented randomly. In each session, the dark filter would be rendered over the left or right eye. 4 sessions were run in counterbalanced order: filter over the left eye, right eye, right eye, then left eye. With four sessions per each of the three observers and 100 trials per session, 1200 trials were collected in total.

During each trial, the dark filter over the left or right eye produced a Pulfrich effect, making the helix appear to rotate in 3 dimensions. The observer nullified the Pulfrich effect by pressing the left and right controller buttons to add 3D motion to the helix stimulus. Observers made inputs with the HTC Vive controller, which has left/right buttons and a trigger. Once the observer decided they had nulled the Pulfrich motion (i.e., the dots appeared to inhabit a fronto-parallel plane), they pressed the trigger button on the controller to advance to the next trial. White noise was flashed between trials to reset the observer’s adaptation state.

### 2.5 Results and Discussion

The main results from Experiment 1 are shown in Figure 3: the Y axis indicates the depth the observer added to the helix to null the 3D rotation (denoted as “rotation adjustment,” in meters); the X axis shows the luminance level of the background. Positive values on the Y axis indicate that the rendered helix motion moves towards the right when the dots are closest to the observer (referred to as “front right” helical motion). Negative values indicate front-left rendered helical motion to null Pulfrich motion. The four symbols represent the conditions of the experiment: black circles indicate black helices with the left eye darkened; gray circles indicate white helices with the left eye darkened; black triangles indicate black helices with the right eye darkened; and gray triangles indicate white helices with the right eye darkened. The error bars show one standard error and are typically smaller than the size of the symbol. The top plot shows the average for all observers; the bottom plots show the results for individual observers.

**Fig. 3.**
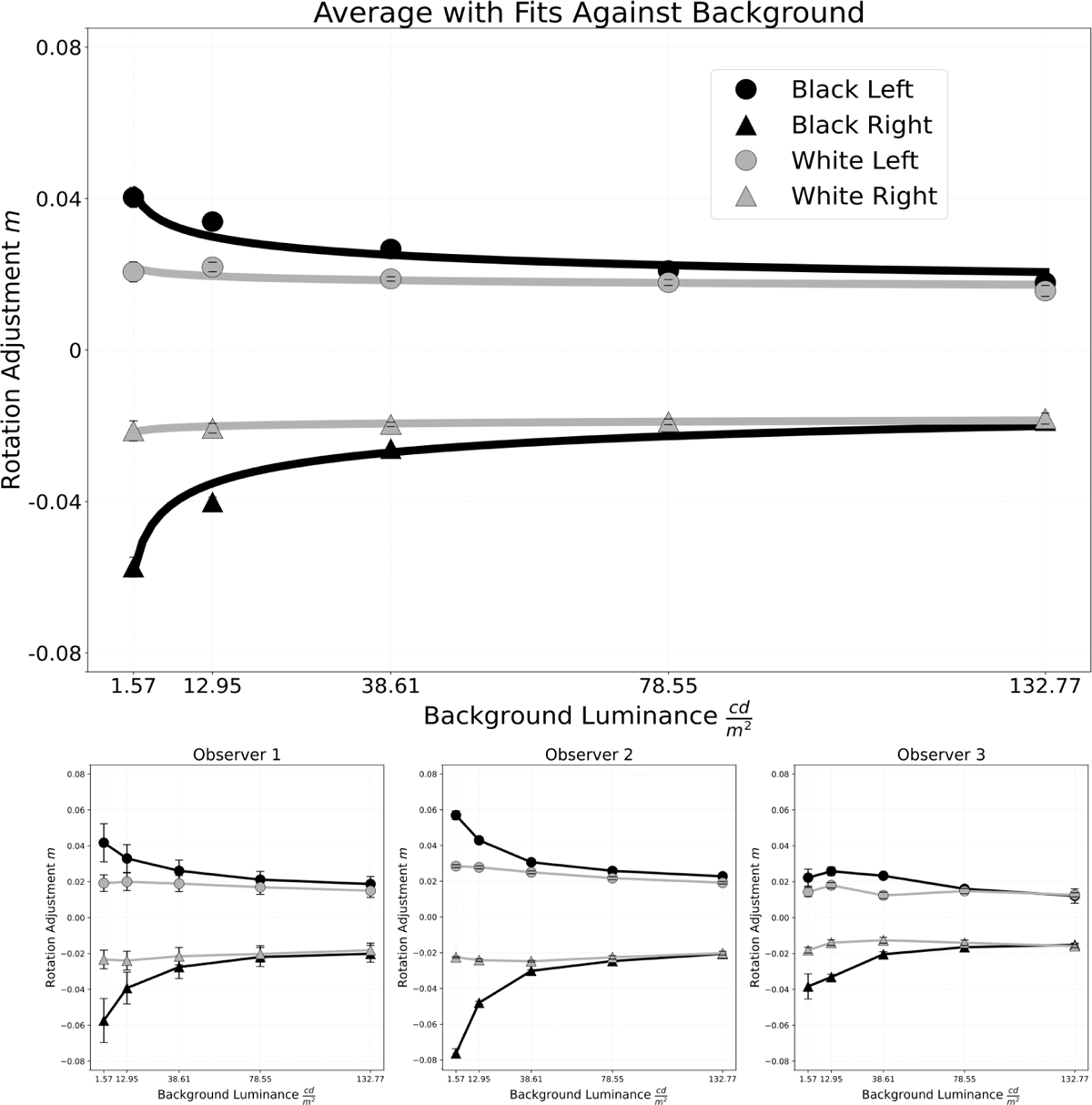
Results from Experiment 1. Observers 1 and 2 are authors AL and AGS respectively, and Observer 3 is the naive observer. The conditions are notated as “color of helix” and “eye being filtered.” For instance, “black left” indicates the session where the helix stimulus was black, and the left eye was being darkened. In “Average with Fits Against Background,” rotation adjustment in meters averaged across all 3 observers for all 4 conditions is plotted as a function of background luminance, alongside standard error bars and power law function fits. Non-visible error bars indicate they are smaller than the plot marker. The bottom 3 plots show individual data for each observer, averaged over each of the 4 sessions.

The main finding was that the magnitude of the Pulfrich effect in virtual reality depends on both overall luminance and local contrast. For all four experimental conditions, decreasing luminance increased the magnitude of rotation adjustment, corroborating a well-known parameter of the Pulfrich effect [24]. However, this effect was more pronounced for the black helix, causing a separation between conditions at the lowest luminance values. At the highest luminance values, the curves of the conditions converged towards each other. These patterns were consistent across the averaged data as well as for each observer’s results. Results will also be interpreted quantitatively through function fits on the averaged data, which are shown in the top plot of Figure 3.

The relationship between perceived depth and background luminance can be described by a power function:

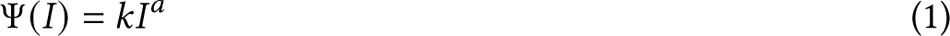

where *I* denotes stimulus intensity (in our case, the luminance of the background), Ψ(*I*) denotes helix rotation necessary to null Pulfrich motion, and *k* and *a* are parameters describing the function behavior. Explicitly, Ψ(*I*) indicates the length of the helix minor axis necessary to null Pulfrich motion. In other words, if *P* is the length of the helix perceived minor axis due to the Pulfrich effect, then *P* + Ψ(*I*) = 0 =⇒ *P* = −Ψ(*I*). Fits were computed with SciPy’s curve-fitting routine, which uses nonlinear least squares to determine function parameters. Function fits were in good agreement with the experimental data from visual assessment.

There were two types of curves: those with steeper slopes (helices with black objects), and those with flatter slopes (helices with white objects). Steep slopes had larger magnitudes of *k* and a parameters (*k* = 0.045 and *a* = −0.16 for black left; *k* = −0.065 and *a* = −0.24 for black right). Flat slopes had smaller magnitudes of k and a parameters (*k* = 0.022 and *a* = −0.05 for white left; *k* = −0.022 and *a* = −0.03 for white right). Flipping which eye is darkened reverses the sign of the k parameter, whereas a was always negative. The parameter differences are notable because they show that changing stimulus reflectance produces perceptual differences that are captured by a power law curve fit. Additionally, there is an apparent asymmetry between the left eye being filtered and the right eye being filtered. Across all observers’ black helix conditions, the right-eye-filtered experiments required approximately 0.2 m greater rotation adjustment magnitude than the left-eye-filtered experiments. However, because the helix rotation method includes motion both towards and away from the filtered eye, the nature of this effect is unclear. It is difficult to make more conclusions about this asymmetry without more testing. Experiment 2 presents a method that allows us to isolate the contributions of motion towards and away from the filtered eye.

Figure 4 plots the average curves as a function of logit-Michelson contrast. The logit function (which is the inverse of the logistic function) is defined as follows:

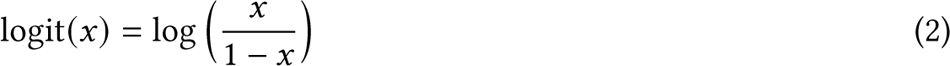

… where 0 *< x <* 1. Michelson contrast is expressed as:

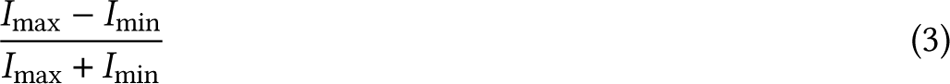

where *I*_max_ and *I*_min_ denote the maximum and minimum luminance of the stimulus, respectively. It should be noted that our stimulus luminance values used for this calculation are determined by mapping machine-unit rendering values of the helix dots to our calibration data. They are therefore only an approximation of the actual luminance of the dots (which depends on position in the virtual environment and is also non-uniform due to specular highlights). Figure 4 shows that contrast only has a strong effect on the black helix data. Furthermore, logit-Michelson contrast appears to linearize the black helix data.

**Fig. 4.**
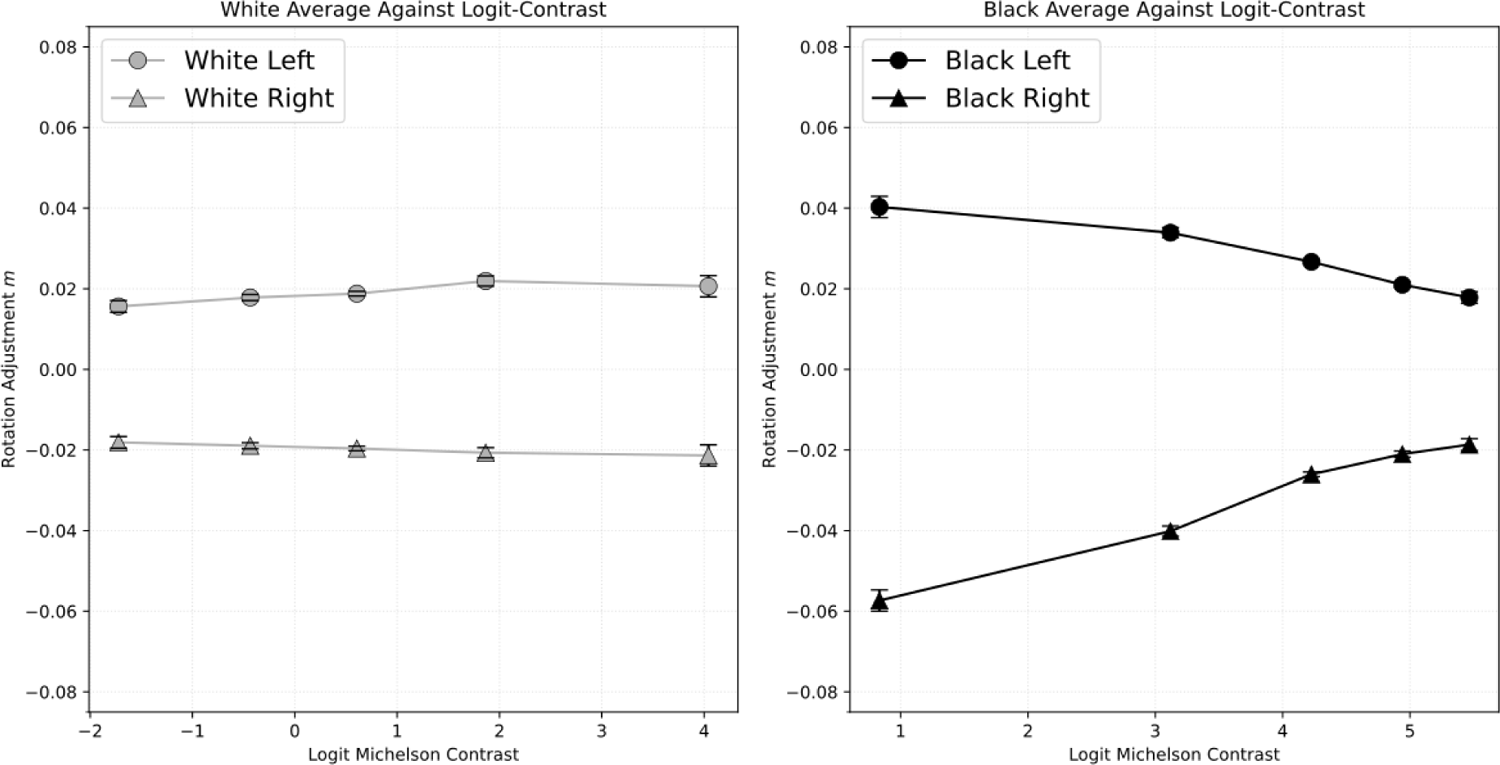
Averaged results from Experiment 1 plotted as a function of logit-Michelson contrast. Michelson contrast was computed with the HMD calibration data, which we note can only produce an approximation. The left plot shows data for the white helix plotted as a function of logit-Michelson contrast; the right plot corresponds to the black helix. Logit-Michelson contrast roughly linearizes the black helix data. The white-helix data, which was already mostly flat, was not robustly affected by the axis transformation.

### 2.6 Illusion Derivable from Experiment 1

Experiment 1 showed that the background creates dramatic differences on the perceived direction of the rotation of black and white helices. The measurements therefore imply that the interplay between luminance and contrast should lead to a novel visual phenomenon in VR. In this illusion, a black helix against a black background appears to rotate towards the right. If the helix is changed from black to white, it will then appear to rotate towards the left instead. The effect also occurs if the background is changed from black to white. That is, a change in the background changes the direction of rotation.

Because the illusion depends on both interocular luminance differences as well as local contrast, it can be thought of as a combination of the Pulfrich and Hess effects. In Figure 3, it is shown that a black helix against a black background needs a minor axis of about 0.04 meters to cancel out the Pulfrich motion when a dark filter is rendered over the left eye. If one instead sets the helix to have a minor axis of 0.03 meters, the helix will continue to rotate towards the left. This is because the motion signal determined by the Pulfrich effect is stronger than the motion signal determined by the helix rotation. If the helix is changed from black to white, the helix motion will overpower the Pulfrich motion because the Pulfrich effect is weaker for high-contrast stimuli than for low-contrast stimuli. As the helix rotation motion to the right is now stronger than the Pulfrich motion to the left, the apparent direction of rotation will switch from left to right. An illustration of the illusion is depicted in Figure 5.

**Fig. 5.**
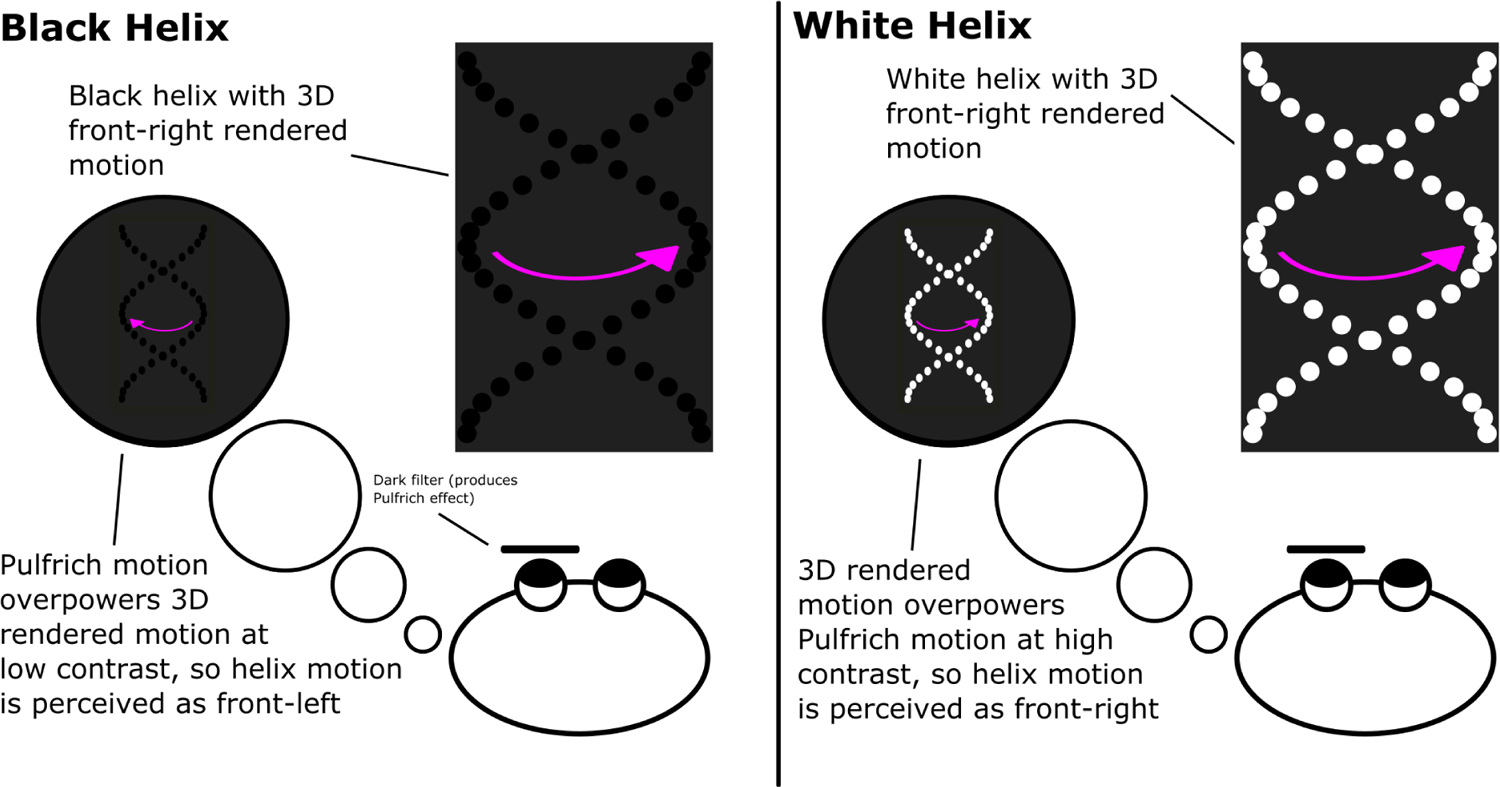
Illustration of the visual illusion derived from our results. The black helix against a dark background produces a stronger Pulfrich effect than the white helix against a dark background. Thus, there is an amount of 3D-rendered motion that will not be strong enough to cancel out the black helix Pulfrich motion but will be strong enough to overpower the white helix Pulfrich motion. Thus, if the 3D-rendered motion of the helix is tuned properly, an effect can be made where a black helix appears to rotate front left, but by changing the color of the helix from black to white, the direction of motion can be changed to front right.

## 3 EXPERIMENT 2: MOVING ARRAY OF DOTS

The Helix configuration of Experiment 1 is a convenient method for measuring depth perception in VR environments. However, the stimulus contains at least two possible confounds related to its motion. First, the sinusoidal back-and-forth motion causes each dot to have several possible perceived depths throughout its trajectory (i.e., a “depth gradient”). Second, the dots move both towards and away from the filtered eye. Here we measure the effect found in Experiment 1 with dots that move at a constant velocity either towards or away from the filtered eye. This new stimulus resolves the issue of the aforementioned confounds as now the dots should only have a single perceived depth and only move towards or away from the filtered eye.

### 3.1 Observers

Three observers participated in this experiment: the two authors and a third naive observer (a different naive observer than in Experiment 1). Observers had normal or corrected vision and no history of colorblindness. Experiments were conducted in accordance with the American University Institutional Review Board. Testing took place on the campus of American University.

### 3.2 Equipment

The equipment and configuration were the same as in Experiment 1.

### 3.3 Stimulus

The stimulus for this experiment was created the same way as in the previous experiment, with everything being generated and rendered with Unity 3D. However, the helix stimulus was replaced with two arrays of dots, as shown in Figure 6. The two arrays of dots were each confined to a single fronto-parallel plane. Dots in the horizontal plane moved laterally across the observer’s field of view at 2.36 m/s. Dots in the vertical plane were stationary. The plane of vertical dots is used as a standard to measure the perceived depth of the plane of horizontally moving dots. Both arrays spanned the entire visual field across their major dimensions. Dots were similar to those in Experiment 1, but now also varied in radius between 0.9 and 2 cm.

**Fig. 6.**
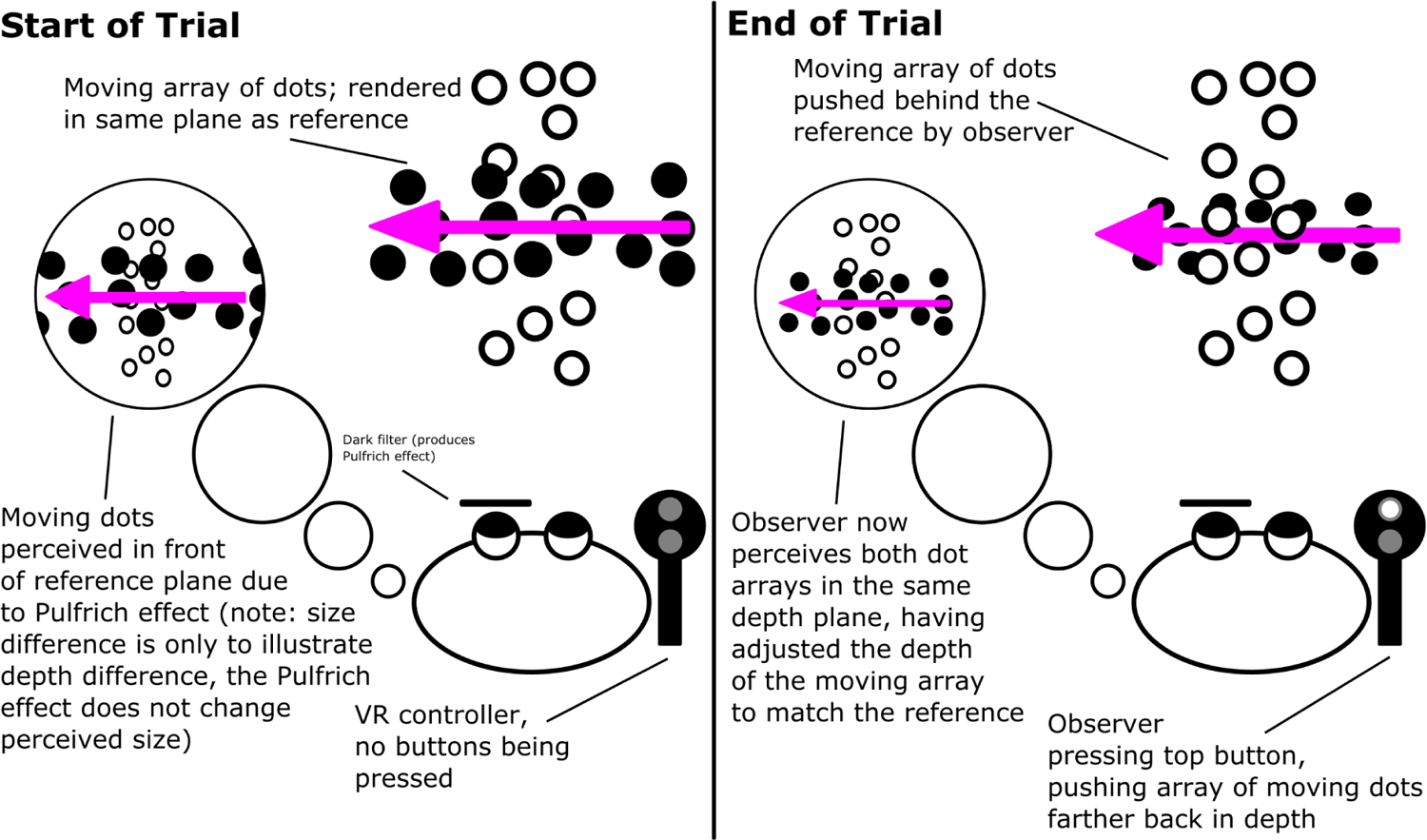
Illustration depicting the task for Experiment 2. There are two planes of dots: vertical stationary dots, which were not affected by the Pulfrich effect, and horizontally moving dots, which were affected by the Pulfrich effect. The task was to adjust the depth plane of horizontal dots so that both the horizontal and vertical dots appeared in the same depth plane.

Each array was always rendered with opposite reflectance values. For instance, if the horizontal array had minimum reflectance (black), then the vertical array had full reflectance (white). During each trial, the background would be rendered at five grayscale RGB intensity values: 0, 0.25, 0.5, 0.75, and 1. These correspond, respectively, to the following luminance values: 0.28, 8.8, 39.7, 93.8, and 180.2 cd/m^2^, as determined via the calibration gamma curve.

### 3.4 Experimental Procedure

The task for this Experiment is depicted in Figure 6. There were four variables in the experiment: which eye was being filtered, the direction of horizontal dot motion, the luminance of the dot arrays, and the luminance of the background. During a given set of sessions for an experiment, either the left or the right eye would be filtered. Each session would then manipulate the color of the dot arrays and the direction of dot motion (left or right).

For each experiment, eight sessions were run for these four conditions (twice per condition) in counterbalanced order. Each session tested depth perception as a function of the five background luminance values. During each session, the filter rendered over the right or left eye would cause the horizontally moving array of dots to appear at a depth. The observer is then tasked with adjusting the depth of the moving array with the Vive controller until it appears to be in the same plane as the stationary array. When the observer believes they have accomplished the task, they press the trigger button on the Vive controller to confirm their selected input. White noise was flashed between each trial upon the Vive trigger press. With 8 trials per condition and background luminance across 4 conditions and 5 background luminance values for the 4 different experiments, 640 trials were collected in total.

### 3.5 Results and Discussion

The results of Experiment 2 are shown in Figure 7. The perceived depth of the moving array of dots is plotted as a function of background luminance as well as logit-Michelson contrast for the averaged data across all observers. For the moving array of dots with motion away from the filtered eye, the strength of the Pulfrich effect increased as overall luminance decreased. Additionally, the magnitude change was stronger for the black helix than for the white helix. However, for motion moving towards the filtered eye, the strength of the Pulfrich effect only slightly increases for the black and white helix as luminance decreases. The top plot shows the averaged data alongside power law fits, computed the same way as in Experiment 1. The bottom plots again show the individual observer plots, noting if the left or right eye was filtered.

**Fig. 7.**
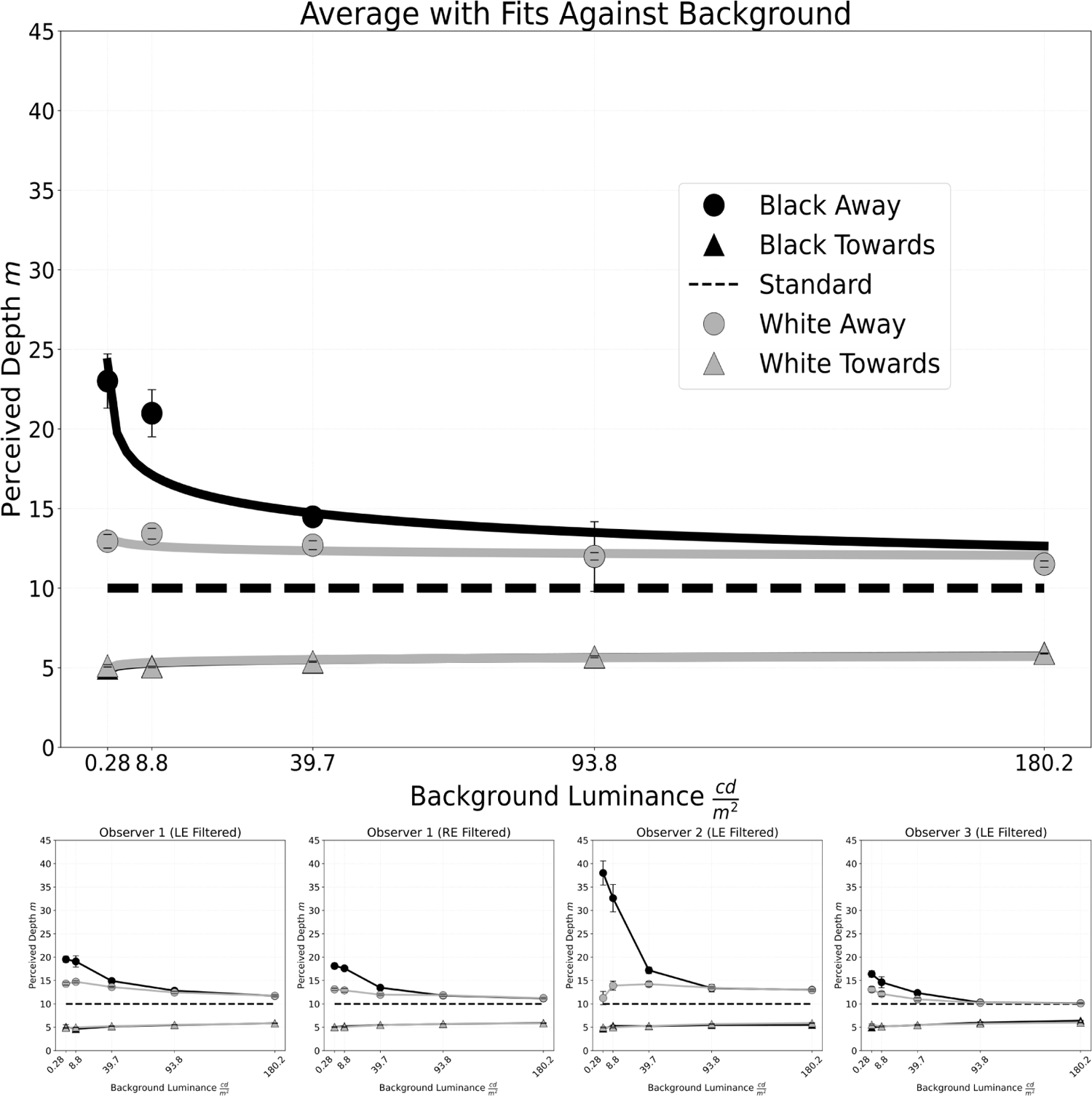
Results for Experiment Two. The conditions are notated as “color of moving array of dots” and “direction of moving array of dots.” For example, the condition “Black Left” indicates the moving array of dots was black and that it was moving towards the left. The titles also indicate which eye is being filtered for the experiment. The dotted line in the middle of each plot is the rendered depth of the standard array, which observers are tasked with adjusting the moving array’s depth to match. Author AL is Observer 1, author AGS is Observer 2, and the naive observer is Observer 3. Due to data collection issues, Observer 1 only had 6 trials for the black right 93.8 cd/m^2^ point.

In Figure 7, the black and white stimulus only separates in the “away” motion condition, indicating that the black/white helix curve separation in Experiment 1 was being driven by motion away from the filtered eye. The difference in depth perception for motion towards and away from the filtered eye could be consistent with the binocular geometry of depth perception. Equivalent disparities correspond to different perceived depths for crossed and uncrossed conditions. Uncrossed disparities, which correspond to the “away” motion in this experiment, produce larger magnitudes of perceived depth. Thus, the effect observed here for the two black curves may actually correspond to the same amount of induced interocular delay, but this effect is not adequately shown due to the binocular geometry of depth-from-disparity perception.

Additionally, there was a wider span of perceived effect strengths across the 3 observers in this experiment than in Experiment 1. Similar to Experiment 1, when the average perceived depth is plotted as a function of logit-Michelson contrast (as shown in Figure 8), the effect strength decreases as contrast increases for the black curve. The effect strength slightly increased as contrast increased for the white curve. In this sense, it can be argued that the motion signal carried by the binocular information is only adequately “captured” for high-luminance and high-contrast stimuli. However, even at lower contrasts, it can be seen that the white and black stimulus at equal contrast levels can have dramatically different perceived depths.

**Fig. 8.**
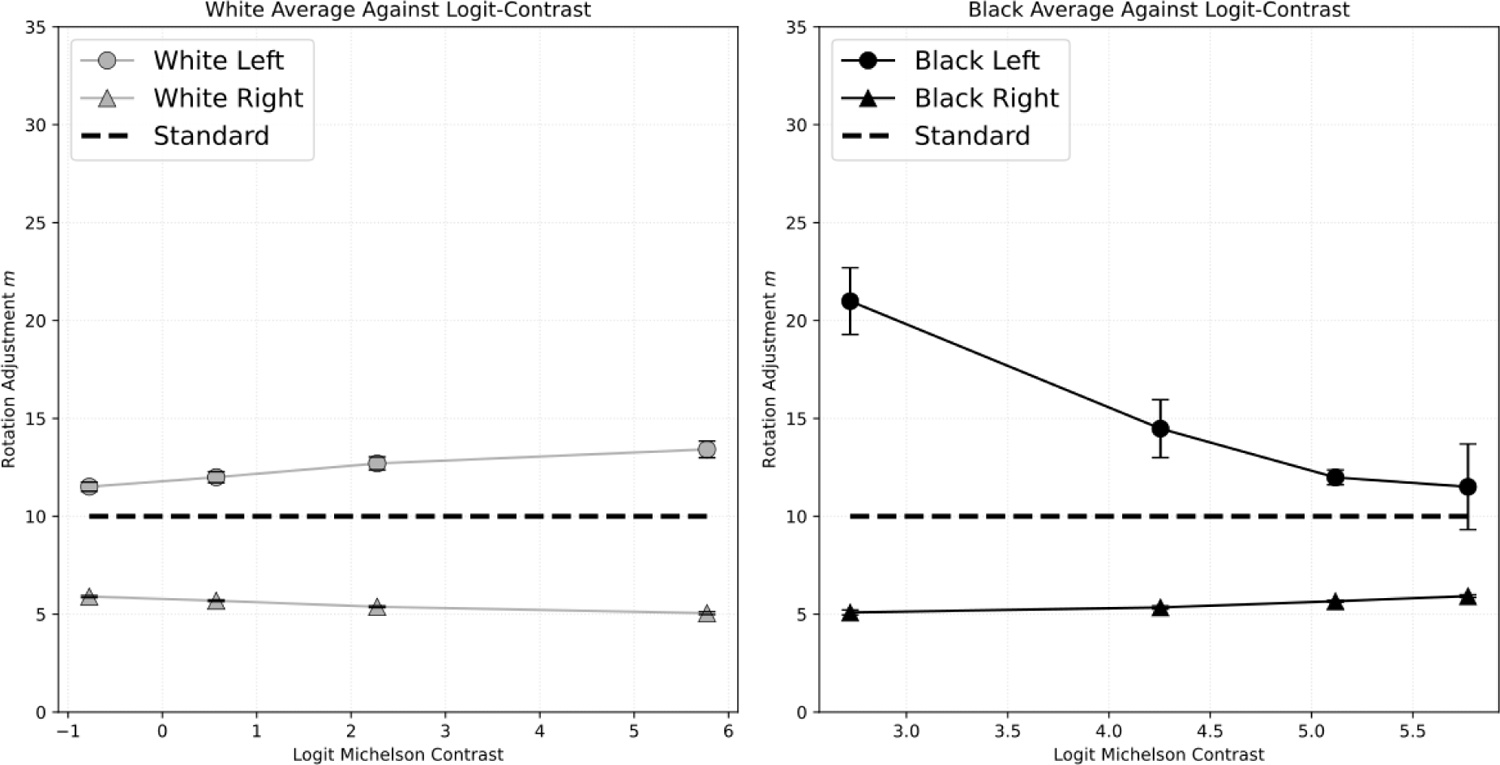
Results for Experiment Two plotted as a function of logit-Michelson contrast. The lowest-luminance background for the black stimulus and the highest-luminance background for the white stimulus are not plotted here because the logarithm of zero is undefined. This data transformation reveals a weak linear relationship for the white curve and a “plateau” effect at higher contrast for the black curve.

## 4 EXPERIMENT 3: HYBRID HELIX MOTION AND BLUR

The previous two experiments documented the relationship between the magnitude of the Pulfrich effect with luminance and contrast in VR environments on HMDs. We have documented that depth perception is affected by binocular luminance differences (Pulfrich effect) and showed that the strength of this effect is mediated by contrast (Hess effect). An additional question can be asked: are these effects influenced by spatial frequency content, and would these effects hold if high spatial frequency content is removed? The question is important at a practical level because visual acuity, and subsequently the spatial frequency content of perceived visual experience, differs from person to person.

We addressed these questions by assessing how optical blur impacts perception of the helix rotation stimulus. This question is related because it illuminates how these two stimulus parameters (luminance contrast and spatial frequency information) are related to motion perception in a stimulus. If jointly increasing spatial frequency and optic blur systematically affect how hybrid motion is perceived, then spatial frequency information in the stimulus may capture the same information as optical blur.

### 4.1 Observers

Three observers participated in this experiment: the two authors and a third naive observer (a different naive observer from that in Experiment 1 and Experiment 2). Observers had normal or corrected vision and no history of colorblindness. Experiments were conducted in accordance with the American University Institutional Review Board. Testing took place on the campus of American University.

### 4.2 Equipment

The experiment was displayed on a 1980 by 1080-pixel 144 Hz ASUS VG248QE monitor. Observers sat 142 cm from the monitor, and the experiment was conducted in a dark room. Observers made inputs using a keyboard seated on their lap. Luminance of the display was calibrated with a Photoresearch Spectroscan 670 spectroradiometer. The maximum luminance for the R, G, and B channels were 31, 101, and 11 cd/m^2^ respectively. The Pulfrich effect was induced with a dark “filter” that observers wore over their glasses, about 0.6 log units. Optic blur was added with 1.5 and 3.0 diopter convex lenses in ophthalmic trial frames adjusted so as to sit in front of the dark filter and observers corrected eyewear.

### 4.3 Stimulus

The stimulus was a green helix in which each “dot” of the helix consists of two orthogonal ellipses that resemble flowers. This stimulus was chosen in order to remain consistent with the experiments in [43]. Each column of the helix contained 12, 18, 24, 30, 36, or 42 dots, with the entire two-column helix containing twice as many dots. The stimulus was 4 degrees wide and 6.6 degrees tall. Each dot of the helix oscillated horizontally to the observer with a frequency of 0.67 Hz. A fixation point is rendered in the center of the helix. In this sense, the stimulus is similar to a version of the VR helix stimulus, with minor variations in order to be more consistent with prior work.

### 4.4 Experimental Procedure

Each of the 5 helices was presented to the observer 5 times in random order per session. During each session, the observer wore the dark filter over their left or right eye, and the trial frame was equipped with 0, 1.5, or 3 diopter lenses. Each of these conditions was run once (2 filter conditions, 3 blur conditions for a total of 6 conditions) in counterbalanced order respective to both the filtered eye and blur: LE, RE, LE, RE, LE, then RE being filtered concurrent with 0, 1.5, 3, 3, 1.5, then 0 diopters of blur. Observers responded to what type of motion they perceived in the helix (left/right motion, or up/down motion) by pressing the respective direction on the arrow pad on their keyboard.

### 4.5 Results and Discussion

The results of Experiment 3 are shown in Figure 9. The top-left plot shows the averaged proportion of trials perceived as up/down versus the number of dots in one column of the helix. The bottom-left plot shows the psychometric functions for the individual observers. The three curves in each of the left plots denote the amount of blur (0, 1.5, and 3.0 diopters). The plots on the right show the threshold number of dots to perceive up or down motion, plotted as a function of optic blur. The top-right plot is for the averaged data, and the bottom-right plots are for the individual observers. Psychometric functions were determined by fitting cumulative Gaussian functions to the data, again using SciPy’s curve-fitting routine which uses nonlinear least squares. Fits were in good agreement with the experimental data. Data is plotted alongside standard error bars for the averaged data. Thresholds were determined from the psychometric fits as the number of dots predicted to produce the proportion of 0.75 up/down responses. The lower row corresponds to individual observer data.

**Fig. 9.**
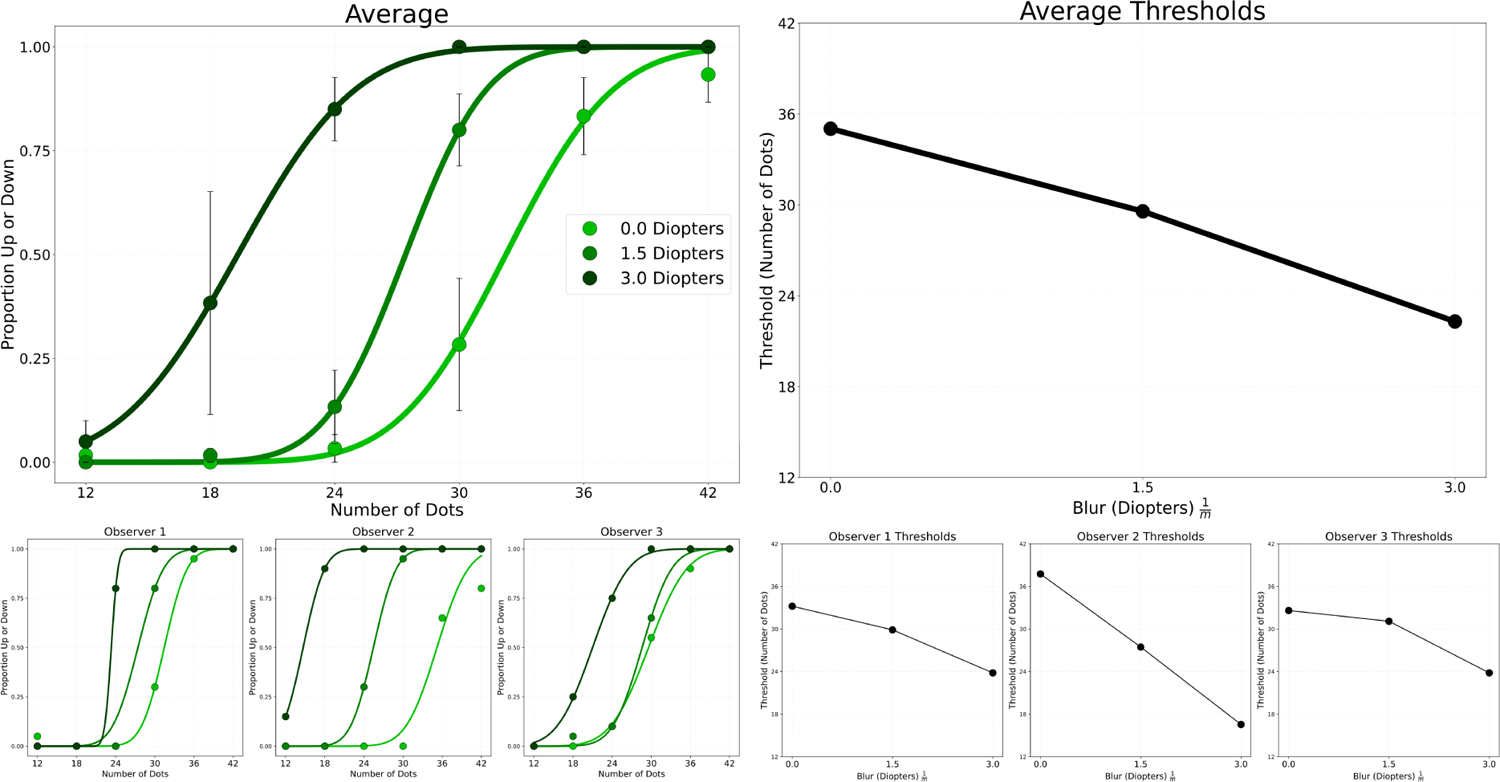
Results for Experiment 3. Observers 1 and 2 are AL and AGS, and Observer 3 is the naive observer. For each observer and the averaged data, the proportion of up/down responses as perceived in the helix stimulus on a flat-panel display is plotted as a function of the number of dots in one column of the helix. The 3 different curves for each of these top four plots denote the amount of optic blur, with the darker curve denoting most blur (3.0 diopters), the next darker curve denoting 1.5 diopters, and the brightest curve denoting no blur. Beneath each plot is the threshold plot for the respective data: the number of dots that produces a proportion of 0.75 up/down responses is plotted as a function of optic blur. In general, the data supports the notion that decreasing dot spacing and increasing optic blur both independently increase the proportion of up/down responses, as elaborated further in the discussion section for this experiment.

There were two main effects. First, an increase in the number of dots in the helix increased the proportion of trials perceived as up/down. That is, perception of the stimulus was driven more by up/down group motion of the dots and less by the Pulfrich effect. Second, increasing the amount of blur shifted this function to the left, indicating that the up/down appearance is easier to produce with few dots in the helix. This general pattern was evident in each observer. As would be expected, the threshold decreases as blur increases across all observers and the averaged data.

These results show that increasing the number of dots and increasing blur both have a similar result on the data, even though increasing the number of dots increases information in the stimulus, whereas adding optic blur decreases information. Increasing either the magnitude of blur or the number of dots leads to the stimulus being perceived as wave-like instead of as a collection of dots. In other words, the stimulus is perceived as a single object instead of as a number of individual features. When this happens, the uncertainty in the stimulus for the location of a single dot decreases, which indicates that the perception of side-to-side motion (i.e., individual dot motion) should become less likely. Likewise, increasing blur also decreases the certainty of each location of a dot but has a smaller effect on the low-spatial frequency information in the stimulus, thereby increasing the proportion of up/down responses.

## 5 GENERAL DISCUSSION

We have introduced a new illusion in which a helix made of black spheres and a helix made of white spheres are perceived to rotate in opposite directions under certain conditions. The illusion was derived by investigating the effect of luminance and contrast on three-dimensional vision in VR. Our finding that increasing background luminance decreases the strength of the Pulfrich effect is consistent with previous studies [24]. However, for the conditions in our first experiment, this effect is much stronger for low-reflectance (black) stimuli than for high-reflectance (white) stimuli, producing two distinct psychometric functions. At the lowest overall background luminance levels, there is a large separation between the strength of the Pulfrich effect for a black helix and a white helix. Therefore, changing the helix from black to white (or equivalently, increasing the contrast of the stimulus against the background) can change the magnitude of the Pulfrich effect. The results indicate an interaction between the Pulfrich and Hess effects, which contribute to processing speed discrepancies due to luminance and contrast, respectively.

The second experiment investigates if Experiment 1’s results were due to motion towards or away from the filtered eye. It was determined that the first experiment’s contrast-dependent effects were driven by motion away from the filtered eye, whereas motion towards the filtered eye showed no contrast dependence. The findings of Experiment 2 are consistent with the mechanics of depth-from-disparity perception.

Experiment 3 examines the effect of dot spacing and blur on hybrid motion perception. It showed that increasing dot count (decreasing dot spacing) and increasing optic blur increase the likelihood of perceiving helical motion as up/down, as opposed to left/right. In this sense, decreasing dot spacing and increasing optic blur both increase dot position uncertainty. This is an interesting finding because increasing optic blur decreases information, and increasing dot count increases information. In spite of this, both operations have the same impact on perception in Experiment 3.

The Pulfrich effect has been widely studied over the course of the past century. It is already well known that the magnitude of the Pulfrich effect increases as light level decreases [24]. Variants of the Pulfrich effect have also been observed for interocular contrast and blur differences [3, 31]. However, to our knowledge, this effect being modulated by contrast between the stimulus and background has not yet been observed. Contrast’s impact on processing speed is already documented [14, 46]. Several other illusions in which changing contrast produces a motion percept have been demonstrated [1, 7, 38]. However, across these examples, the information provided by contrast is noted to be a “weak” motion signal. In other words, the motion illusions produced by changing contrast for a stationary object would be overridden if that object were to physically move. In our effects, however, contrast can completely reverse the trajectory of a moving object. This seems to indicate that in certain circumstances, contrast can be a strong motion and depth cue rather than a weak one.

To quickly and informally confirm that the effects observed across Experiments 1 and 2 were not due to hardware or software issues, author AL conducted two tests. First, AL tested to see if the observed visual illusion worked with a dark filter placed over the HMD screens and observed that the illusion still worked the same with a physical filter as with the rendered filter. Additionally, AL created a demo version of the experiment that ran on an Oculus Quest headset and again observed that the illusion worked the same as it did on the HTC Vive. Though informal observations such as these are not equivalent to proper trial-randomized experiment reproduction, they at least provide some assurance that the results of these experiments are legitimate. It is possible, however, that the results are due to the rendering of the stimuli itself in Unity (with specular highlights and complex lighting cues) and not due to anything unique to perception in VR HMDs.

However, it is also possible that these results are unique to VR HMDs, as there are many distinctions between VR systems and natural viewing or haploscopes commonly used in vision research. One important distinction between VR HMDs and natural viewing is the vergence-accommodation conflict [5, 20]. In the vergence-accommodation conflict, the oculomotor depth cues of vergence (angle between the eyes) and accommodation (optical power of the crystalline lens behind the eye) provide different information about the depth of a virtual object. Additionally, VR HMDs are often referred to as “near-eye displays,” because the displays are mere centimeters from your eyes, despite the fact that the virtual images produced by the displays can be at varying distances [22]. VR headsets also have poor resolution relative to human visual acuity [25]. Our calibration method may have also impacted the results, as it is now known that properly calibrating a VR headset running Unity applications is more involved than it is for other setups [28]. The ability for viewers to move their head freely in a VR environment is also different from classical psychophysical experiments, which often keep stimuli at a fixed point on the retina. We do not claim any differences between perception in VR and the natural world causally explain the effects in this paper. However, we do claim that reproducing old vision science experiments in VR and searching for new results could be of importance because of the discrepancies our experiments have identified.

The tradition of illusion research is motivated by finding compelling visual stimuli whether or not they immediately elucidate new underlying mechanisms of neural processing [19]. In spite of this, these stimuli often lead to the discovery of new mechanisms or refine understanding of old mechanisms. In this sense, reproducing or creating visual illusions in VR may have the power to generalize our understanding of old illusions and mechanisms. Here, we investigated the Pulfrich effect in VR and found a unique contrast effect on depth and motion perception. Although this finding is novel, the Pulfrich effect in virtual reality in itself is still compelling, as is how visually impressive the nulling procedure is. This is not the first study on “illusions” in VR: previous studies have already investigated color awareness [6], contrast asynchronies [33], and the Ebbinghaus illusion [47], as well as “the rubber hand illusion” [15, 34] However, we do believe that reproducing older illusions such as the Pulfrich effect is of high value. Other classic illusions that could be investigated in VR include Kanizsa and checker shadow illusions.

It is common in vision science psychophysics experiments to use a small-N design with a large number of trials per observer [11, 45]. Historically, seminal Pulfrich effect psychophysics papers are low-n studies featuring 2 or 3 observers [24, 27] and effects like these seldom have problems with replication. Modern studies often employ around 5-10 observers [3, 26]. Our study offers a compromise between earlier and modern studies: each experiment is a low-n (3) observer study, but each experiment in this paper utilizes a distinct naive observer. With the two authors in each experiment and 3 different naive observers across all 3 experiments, this paper uses a total of 5 observers. It is conceivable that larger numbers of observers could change the trends of these effects. We think this is unlikely as our variations of the Pulfrich effect have been used in both public and lab demonstrations, and the phenomenon reported here seems to be very robust.

This study investigated only black and white stimuli in virtual reality. The question of what curves may we expect to see for gray stimuli is therefore left open. Figure 10 presents two plausible theories: that psychometric functions for gray stimuli will be between the curves for the black and white helices, or that the curves for gray helices will converge (or be “attracted”) to the black and white psychometric functions. The first theory is consistent with continuous representations of luminance, whereas the second theory is consistent with a discrete mechanism determining if luminance is above or below a particular threshold (i.e., luminance “capture” of information in the stimulus.) Testing the perception of gray stimuli in VR under the Pulfrich effect could therefore be an important future experiment.

**Fig. 10.**
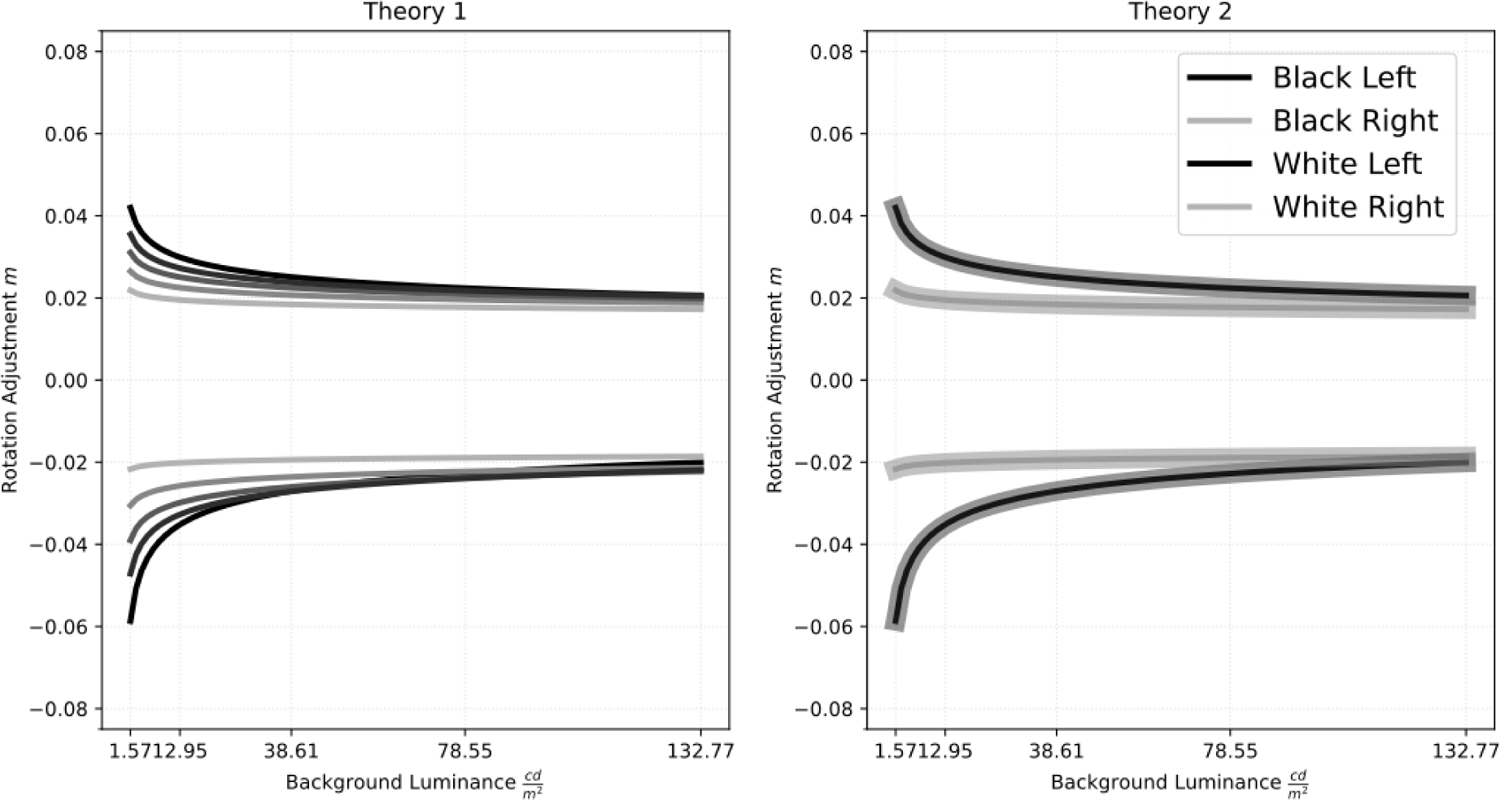
Two theories for what psychometric functions may look like for gray helices. In Theory 1, gray helices “span” the area bounded by the curves determined by the black and white stimuli. In the second theory, gray helices are instead “attracted” to the curves for the black and white helices, in a way consistent with contrast “capturing” either the binocular disparity motion or the Pulfrich effect motion.

Lastly, the experiments address questions related to “background-independent constancy” [10, 50]. There are two different types of color constancy [2]: the type most frequently discussed concerns the constant appearance of a surface under different types of illumination. “Background-independent constancy” refers to the relatively constant appearance of an object viewed against different natural backgrounds. This type of constancy may seem odd because, in the laboratory, it is possible to create demonstrations where the background plays an important role in how foreground objects are perceived. For example, a gray square’s perceived brightness is modulated by the luminance of the background it is presented against in an experimental setup. A real-world equivalent (such as a square piece of paper) most often appears similar whether it’s viewed against a backdrop of a forest or a featureless wall. One argument for the distinction is that experimental setups remove some natural variables like binocular disparity and the ability to move one’s head freely. Additionally, the changes in perceived brightness induced by different luminances of a uniform background can hardly be denoted as striking. In our experiment, not only are some naturalistic features present due to the use of VR, but the background plays a salient role in perception by altering the perceived trajectory of a moving object. As such, by pitting binocular depth cues from the Pulfrich effect against background-based pictorial cues from the Hess effect, it may be possible to create situations where the background plays a more salient role in perception even in experiments more naturalistic than ours.

## 6 CONCLUSION

We have used the conflict between Pulfrich and Hess phenomena to investigate the interaction between luminance and contrast and their effects on depth perception in a virtual reality (VR) environment. We show that low-reflectance stimuli greatly increase the strength of the Pulfrich effect, whereas high-reflectance stimuli slightly increase the strength. To our knowledge, the darker stimulus increasing the strength of the Pulfrich effect is unreported and may be unique to the viewing conditions of VR displays. The results show that the helix rotation nulling method is a novel and extremely powerful way to study the magnitude of the Pulfrich effect. Additionally, this result can be used to create a new type of highly compelling visual phenomenon, where the direction of motion of the helix changes simply by changing the luminance (i.e., contrast) of the stimulus. We further investigate this effect with a second experiment that disentangles the types of motion in the first experiment, and postulate the existence of an effect related to dot spacing with a non-VR experiment studying the effect of motion perception in the helix as it relates to dot spacing and blur.

The results of these experiments may prove useful when thinking about how to design VR environments, which typically are concerned only with depth from binocular disparity. Our results highlight the need to study old effects in VR. Not only do VR technologies allow us to study laboratory effects in more naturalistic scenarios, but, given the increasing prevalence of the headsets and their unusual way of integrating with the human visual system (as elaborated upon in the discussion), observers could experience any number of surprising phenomena. As such, taking old illusions and studying them in VR opens up a window to understand how perception in these environments may differ from traditional laboratory setups and may even elucidate important properties of perception.

## ACKNOWLEDGMENTS

This work was supported by a fellowship from the NASA DC Space Grant Consortium.

## REFERENCES

[1] Stuart M Anstis and Brian J Rogers. 1986. Illusory Continuous Motion from Oscillating Positive-Negative Patterns: Implications for Motion Perception. Perception 15, 5 (Oct. 1986), 627–640. 10.1068/p150627 Publisher: SAGE Publications Ltd STM.

[2] Richard O. Brown. 2003. Backgrounds and Illuminants: The Yin and Yang of Colour Constancy. In Colour Perception: Mind and the physical world, Rainer Mausfeld and Dieter Heyer (Eds.). Oxford University Press, 0. 10.1093/acprof:oso/9780198505006.003.0008

[3] Johannes Burge, Victor Rodriguez-Lopez, and Carlos Dorronsoro. 2019. Monovision and the Misperception of Motion. Current Biology 29, 15 (Aug. 2019), 2586–2592.e4. 10.1016/j.cub.2019.06.070

[4] Benjamin M. Chin and Johannes Burge. 2022. Perceptual consequences of interocular differences in the duration of temporal integration. Journal of Vision 22, 12 (Nov. 2022), 12. 10.1167/jov.22.12.12

[5] Steven A. Cholewiak, Zeynep Başgöze, Ozan Cakmakci, David M. Hoffman, and Emily A. Cooper. 2020. A perceptual eyebox for near-eye displays. Optics Express 28, 25 (Dec. 2020), 38008–38028. 10.1364/OE.408404 Publisher: Optica Publishing Group.

[6] Michael A. Cohen, Thomas L. Botch, and Caroline E. Robertson. 2020. The limits of color awareness during active, real-world vision. Proceedings of the National Academy of Sciences 117, 24 (June 2020), 13821–13827. 10.1073/pnas.1922294117 Publisher: Proceedings of the National Academy of Sciences.

[7] Oliver J. Flynn and Arthur G. Shapiro. 2018. The Perpetual Diamond: Contrast Reversals Along Thin Edges Create the Appearance of Motion in Objects. i-Perception 9, 6 (Nov. 2018), 2041669518815708. 10.1177/2041669518815708 Publisher: SAGE Publications.

[8] Jacqueline M. Fulvio and Bas Rokers. 2017. Use of cues in virtual reality depends on visual feedback. Scientific Reports 7, 1 (Nov. 2017), 16009. 10.1038/s41598-017-16161-3 Number: 1 Publisher: Nature Publishing Group.

[9] Raquel Gil Rodríguez, Florian Bayer, Matteo Toscani, Dar’ya Guarnera, Giuseppe Claudio Guarnera, and Karl R. Gegenfurtner. 2021. Colour Calibration of a Head Mounted Display for Colour Vision Research Using Virtual Reality. SN Computer Science 3, 1 (Oct. 2021), 22. 10.1007/s42979-021-00855-7

[10] Alan Gilchrist. 2006. Seeing Black and White. Oxford University Press. Google-Books-ID: ChriBwAAQBAJ.

[11] James E. Graham, Amol M. Karmarkar, and Kenneth J. Ottenbacher. 2012. Small Sample Research Designs for Evidence-based Rehabilitation: Issues and Methods. Archives of physical medicine and rehabilitation 93, 8 Suppl (Aug. 2012), S111–S116. 10.1016/j.apmr.2011.12.017

[12] Rick Gurnsey and Mathieu Biard. 2012. Eccentricity dependence of the curveball illusion. Canadian Journal of Experimental Psychology 66, 2 (2012), 144–152. 10.1037/a0026989 Place: US Publisher: Educational Publishing Foundation.

[13] Amanda J. Haskins, Jeff Mentch, Thomas L. Botch, and Caroline E. Robertson. 2020. Active vision in immersive, 360° real-world environments. Scientific Reports 10, 1 (Aug. 2020), 14304. 10.1038/s41598-020-71125-4 Number: 1 Publisher: Nature Publishing Group.

[14] Carl von Hess. 1904. Untersuchungen über den Erregungsvorgang im Sehorgan bei kurz-und bei längerdauernder Reizung. Archiv für die gesamte Physiologie des Menschen und der Tiere 101, 5 (1904), 226–262.

[15] Wijnand A IJsselsteijn, Yvonne A. W de Kort, and Antal Haans. 2006. Is This My Hand I See Before Me? The Rubber Hand Illusion in Reality, Virtual Reality, and Mixed Reality. Presence: Teleoperators and Virtual Environments 15, 4 (Aug. 2006), 455–464. 10.1162/pres.15.4.455

[16] E. L. Isenstein, T. Waz, A. LoPrete, Y. Hernandez, E. J. Knight, A. Busza, and D. Tadin. 2022. Rapid assessment of hand reaching using virtual reality and application in cerebellar stroke. PLOS ONE 17, 9 (Sept. 2022), e0275220. 10.1371/journal.pone.0275220 Publisher: Public Library of Science.

[17] Jonathan W. Kelly, Lucia A. Cherep, Brenna Klesel, Zachary D. Siegel, and Seth George. 2018. Comparison of Two Methods for Improving Distance Perception in Virtual Reality. ACM Transactions on Applied Perception 15, 2 (March 2018), 11:1–11:11. 10.1145/3165285

[18] Frederick A. A. Kingdom, David R. Simmons, and Stéphane Rainville. 1999. On the apparent collapse of stereopsis in random-dot-stereograms at isoluminance. Vision Research 39, 12 (June 1999), 2127–2141. 10.1016/S0042-6989(98)00257-0

[19] Akiyoshi Kitaoka, Takahiro Kawabe, and Yuki Yamada. 2020. Introducing the Journal of Illusion. Journal of Illusion 1 (Oct. 2020). 10.47691/joi.v1.5591

[20] Gregory Kramida. 2016. Resolving the Vergence-Accommodation Conflict in Head-Mounted Displays. IEEE Transactions on Visualization and Computer Graphics 22, 7 (July 2016), 1912–1931. 10.1109/TVCG.2015.2473855 Conference Name: IEEE Transactions on Visualization and Computer Graphics.

[21] Oh-Sang Kwon, Duje Tadin, and David C. Knill. 2015. Unifying account of visual motion and position perception. Proceedings of the National Academy of Sciences 112, 26 (June 2015), 8142–8147. 10.1073/pnas.1500361112 Publisher: Proceedings of the National Academy of Sciences.

[22] Douglas Lanman and David Luebke. 2013. Near-eye light field displays. ACM Transactions on Graphics 32, 6 (Nov. 2013), 220:1–220:10. 10.1145/2508363.2508366

[23] Matteo Lisi and Patrick Cavanagh. 2015. Dissociation between the Perceptual and Saccadic Localization of Moving Objects. Current biology: CB 25, 19 (Oct. 2015), 2535–2540. 10.1016/j.cub.2015.08.021

[24] Alfred Lit. 1949. The Magnitude of the Pulfrich Stereophenomenon as a Function of Binocular Differences of Intensity at Various Levels of Illumination. The American Journal of Psychology 62, 2 (1949), 159–181. 10.2307/1418457 Publisher: University of Illinois Press.

[25] Marissa Howard Lynn, Gang Luo, Matteo Tomasi, Shrinivas Pundlik, and Kevin E. Houston. 2020. Measuring Virtual Reality Headset Resolution and Field of View: Implications for Vision Care Applications. Optometry and Vision Science 97, 8 (Aug. 2020), 573–582. 10.1097/OPX.0000000000001541

[26] Seung Hyun Min, Alexandre Reynaud, and Robert F. Hess. 2020. Interocular Differences in Spatial Frequency Influence the Pulfrich Effect. Vision 4, 1 (March 2020), 20. 10.3390/vision4010020 Number: 1 Publisher: Multidisciplinary Digital Publishing Institute.

[27] Michael J Morgan and Peter Thompson. 1975. Apparent Motion and the Pulfrich Effect. Perception 4, 1 (March 1975), 3–18. 10.1068/p040003 Publisher: SAGE Publications Ltd STM.

[28] Richard F. Murray, Khushbu Y. Patel, and Emma S. Wiedenmann. 2022. Luminance calibration of virtual reality displays in Unity. Journal of Vision 22, 13 (Dec. 2022), 1. 10.1167/jov.22.13.1

[29] Aude Oliva, Antonio Torralba, and Philippe G. Schyns. 2006. Hybrid images. ACM Transactions on Graphics 25, 3 (July 2006), 527–532. 10.1145/1141911.1141919

[30] Carl Pulfrich. 1922. Die Stereoskopie im Dienste der isochromen und heterochromen Photometrie. Die Naturewissenschaften 10, 35 (1922), 553–564.

[31] Alexandre Reynaud and Robert F. Hess. 2017. Interocular contrast difference drives illusory 3D percept. Scientific Reports 7, 1 (July 2017), 5587. 10.1038/s41598-017-06151-w Number: 1 Publisher: Nature Publishing Group.

[32] Bernhard Riecke, Joerg schulte pelkum, Marios Avraamides, and Heinrich Bülthoff. 2004. Enhancing the visually induced self-motion illusion (vection) under natural viewing conditions in virtual reality. Proceedings of Seventh Annual Workshop Presence 2004 (Jan. 2004).

[33] Alex Rose-Henig and Arthur G. Shapiro. 2014. Contrast–contrast asynchronies. JOSA A 31, 4 (April 2014), A232–A238. 10.1364/JOSAA.31.00A232 Publisher: Optica Publishing Group.

[34] Anca Salagean, Jacob Hadnett-Hunter, Daniel J. Finnegan, Alexandra A. De Sousa, and Michael J. Proulx. 2022. A Virtual Reality Application of the Rubber Hand Illusion Induced by Ultrasonic Mid-air Haptic Stimulation. ACM Transactions on Applied Perception 19, 1 (Jan. 2022), 3:1–3:19. 10.1145/3487563

[35] Peter Scarfe and Andrew Glennerster. 2019. The Science Behind Virtual Reality Displays. Annual Review of Vision Science 5, 1 (2019), 529–547. 10.1146/annurev-vision-091718-014942.

[36] Lauren V. Scharff and Wilson S. Geisler. 1992. Stereopsis at isoluminance in the absence of chromatic aberrations. JOSA A 9, 6 (June 1992), 868–876. 10.1364/JOSAA.9.000868 Publisher: Optica Publishing Group.

[37] Arthur Shapiro. 2021. Hybrid motion illusions as examples of perceptual conflict. Journal of Illusion 2 (Sept. 2021). 10.47691/joi.v2.7084

[38] Arthur Shapiro and Emily Knight. 2008. Spatial and temporal influences on the contrast gauge. Vision Research 48, 26 (Nov. 2008), 2642–2648. 10.1016/j.visres.2008.06.027

[39] Arthur Shapiro, Zhong-Lin Lu, Chang-Bing Huang, Emily Knight, and Robert Ennis. 2010. Transitions between Central and Peripheral Vision Create Spatial/Temporal Distortions: A Hypothesis Concerning the Perceived Break of the Curveball. PLOS ONE 5, 10 (Oct. 2010), e13296. 10.1371/journal.pone.0013296 Publisher: Public Library of Science.

[40] Arthur G. Shapiro. 2008. Separating color from color contrast. Journal of Vision 8, 1 (Jan. 2008), 8. 10.1167/8.1.8

[41] Arthur G. Shapiro, Justin P. Charles, and Mallory Shear-Heyman. 2005. Visual illusions based on single-field contrast asynchronies. Journal of Vision 5, 10 (Nov. 2005), 2. 10.1167/5.10.2

[42] Arthur G Shapiro and Laysa Hedjar. 2019. Color illusion as a spatial binding problem. Current Opinion in Behavioral Sciences 30 (Dec. 2019), 149–155. 10.1016/j.cobeha.2019.08.004

[43] Arthur G. Shapiro and Anthony LoPrete. 2020. Helix rotation: luminance contrast controls the shift from two-dimensional to three-dimensional perception. JOSA A 37, 4 (April 2020), A262–A270. 10.1364/JOSAA.382373 Publisher: Optica Publishing Group.

[44] Vincent Sitzmann, Ana Serrano, Amy Pavel, Maneesh Agrawala, Diego Gutierrez, Belen Masia, and Gordon Wetzstein. 2018. Saliency in VR: How Do People Explore Virtual Environments? IEEE Transactions on Visualization and Computer Graphics 24, 4 (April 2018), 1633–1642. 10.1109/TVCG.2018.2793599 Conference Name: IEEE Transactions on Visualization and Computer Graphics.

[45] Philip L. Smith and Daniel R. Little. 2018. Small is beautiful: In defense of the small-N design. Psychonomic Bulletin & Review 25, 6 (Dec. 2018), 2083–2101. 10.3758/s13423-018-1451-8

[46] Peter Thompson and Stuart Anstis. 2005. Retracing our footsteps: A revised theory of the footsteps illusion. Journal of Vision 5, 8 (Sept. 2005), 929. 10.1167/5.8.929

[47] Russell Todd, Qin Zhu, and Amy Banić. 2021. Temporal Availability of Ebbinghaus Illusions on Perceiving and Interacting with 3D Objects in a Contextual Virtual Environment. In 2021 IEEE Virtual Reality and 3D User Interfaces (VR). 817–825. 10.1109/VR50410.2021.00109 ISSN: 2642-5254.

[48] Anne Treisman. 1996. The binding problem. Current Opinion in Neurobiology 6, 2 (April 1996), 171–178. 10.1016/S0959-4388(96)80070-5

[49] P. U. Tse and P. J. Hsieh. 2006. The infinite regress illusion reveals faulty integration of local and global motion signals. Vision Research 46, 22 (Oct. 2006), 3881–3885. 10.1016/j.visres.2006.06.010

[50] Paul Whittle. 1994. The psychophysics of contrast brightness. In Lightness, brightness, and transparency. Lawrence Erlbaum Associates, Inc, Hillsdale, NJ, US, 35–110.

[51] Laurie M. Wilcox, Robert S. Allison, Samuel Elfassy, and Cynthia Grelik. 2006. Personal space in virtual reality. ACM Transactions on Applied Perception 3, 4 (Oct. 2006), 412–428. 10.1145/1190036.1190041

[52] John Michael Williams and Alfred Lit. 1983. Luminance-dependent visual latency for the hess effect, the pulfrich effect, and simple reaction time. Vision Research 23, 2 (Jan. 1983), 171–179. 10.1016/0042-6989(83)90140-2

